# Transient cortical Beta-frequency oscillations associated with contextual novelty in high density mouse EEG

**DOI:** 10.1101/2024.07.09.602651

**Authors:** Callum Walsh, Luke Tait, Maria Garcia Garrido, Jonathan T. Brown, Thomas Ridler

## Abstract

Beta-frequency oscillations (20-30 Hz) are prominent in both human and rodent electroencephalogram (EEG) recordings. Discrete epochs of beta (or Beta2) oscillations are prevalent in the hippocampus and other brain areas during exploration of novel environments. However, little is known about the spatial distribution and temporal relationships of beta oscillations across the cortex in response to novelty. To investigate this, mice fitted with 30-channel EEG-style multi-electrode arrays underwent a single recording session in a novel environment. While changes to spectral properties of cortical oscillations were minimal, there was a profound increase in the rate of beta bursts during the initial part of the recording session, when the environment was most novel. This was true across the cortex but most notable in recording channels situated above the retrosplenial cortex. Additionally, novelty was associated with greater connectivity between retrosplenial areas and the rest of the cortex, specifically in the beta frequency range. However, it was also found that the cortex in general, is highly modulated by environmental novelty. This data further suggests the retrosplenial cortex is an important hub for distinguishing environmental context and highlights the diversity of functions for beta oscillations across the brain, which can be observed using high-density EEG.

## Introduction

The ability to distinguish environmental novelty is vitally important for providing both appropriate behavioural responses and for the accurate encoding of new representations of environments. In rodents, the relationship between beta (20-30 Hz) oscillations (sometimes referred to as Beta2) and novelty has been known for some time^1^. Discrete epochs of oscillations at this beta frequency , or beta bursts, are prominent in the hippocampus during the first minutes of exploration in novel environments^1,2^ and when interacting with novel objects, suggesting a potential role in the encoding of novel information^2–4^. The incidence of beta bursting decreases dramatically as environments or stimuli become familiar. More recently, several other cortical regions have been shown to participate in these bursts. Beta bursts have been detected in parietal and mid-frontal cortices and are highly coherent with those in the hippocampus^2,5^. In addition, beta bursting has been detected in the retrosplenial cortex, where they increase in prevalence during contextual novelty and are associated with transient increases in neuronal spiking^6^, further highlighting this region as an important area for context discrimination. Activation, and later reactivation, of neuronal ensembles during beta bursts may facilitate the creation and recall of cortical representations of environments ^7–9^.

This said, relatively little is known about the spatial distribution of beta oscillations across the entire cortical surface in response to environmental novelty. Beta bursting has been previously described in a range of brain regions and is associated with a wide variety of behaviours^10–16^. Spontaneous beta bursts have been identified in the somatosensory cortex and frontal cortex in humans, rodents, and non-human primates^12^, and pre-stimulus beta bursting in the somatosensory cortex of mice and humans is negatively correlated with tactile stimulus detection^17^. In motor cortex in humans, beta bursts were associated with the termination of movement^11^, and in another study the timing of motor cortex beta bursts was associated with the timing of movement initiation, and errors were associated with delayed or reduced beta bursting^13^. The role of beta bursting may therefore vary dramatically depending on its location within the brain^11^ and while beta oscillations may be prevalent across the cortex, their relationship with novelty is likely to be confined to specific areas.

We therefore aimed to investigate novelty-associated beta bursting across the cortex and determine the temporal relationships between cortical beta oscillations. To investigate beta bursting across the cortex, mice were fitted with EEG-style multi-electrode arrays covering the dorsal cortical surface and local field potentials were recorded while mice explored a novel environment. We demonstrate that spontaneous beta bursting occurs across the cortex, but novelty-associated beta bursting appears preferentially localised around the retrosplenial cortex.

## Results

To investigate the response of the cortical EEG to environmental novelty we implanted 30 channel EEG-style electrode arrays onto the skull surface and exposed animals to a completely novel recording environment (Fig. 1A-D). Over the course of the 15-minute recording session, mice have been shown to quickly become familiar with this arena, with the initial 1-2 minutes of exploration having previously been associated with enhanced beta oscillations in a variety of brain regions^6,18^. To compare between the novel part of the session and the familiar part of the session we split our recordings into the first minute (initial/novel) and last 10 minutes (final/familiar) of the recording.

**Figure 1:**
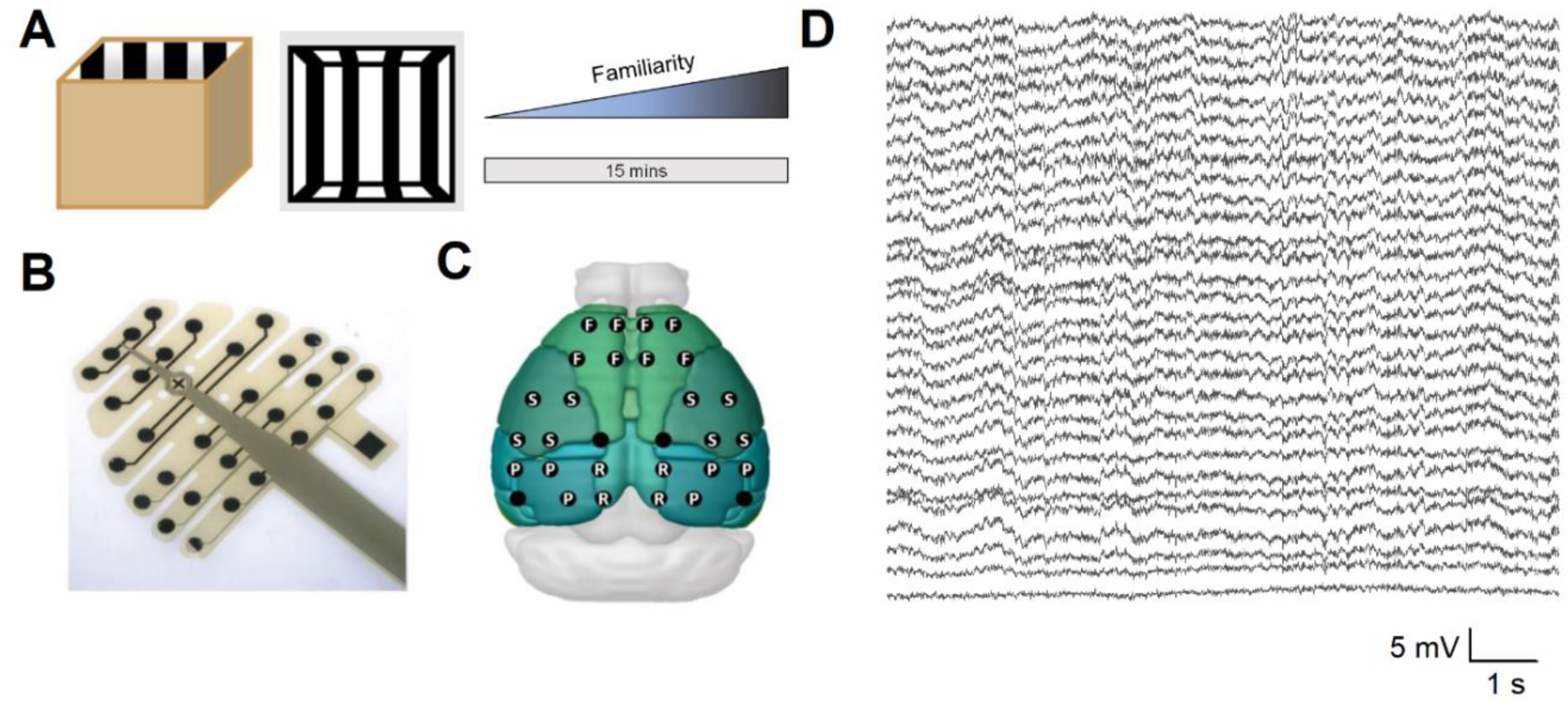
High density EEG recording in novel environment. **A)** Schematic of 1m x 1m recording arena (left) and schematic to represent increasing familiarity with the environment over the recording session. **B)** Picture of electrode array from NeuroNexus. Circles: electrode locations, Cross: Bregma, Square: Reference channel. **C)** Superimposed electrode locations of EEG channels over the cortical surface of the mouse brain. For some analyses, the channels were grouped into four approximate cortical areas and averaged. These were: F = Frontal, S = Somatosensory, P = Parietal, and R = Retrosplenial. 4 electrodes were not classified, as they did not fit clearly into any of these areas (empty dots). **D)** Example traces from all 30 electrodes during novel recording session (scale bar: 5 mV, 1 s). We first performed spectral analysis, comparing the spectral power in the Theta (5-12 Hz), Beta (20-30 Hz) and Gamma (30-100 Hz) frequency ranges between the initial and final parts of the session. For all frequency bands tested there were topographical variations in the power of oscillatory activity, typically in the form of higher power oscillations in the frontal recording area; however, oscillatory power did not appear to differ significantly between novel (initial) and familiar (final) timepoints (Fig. 2A-C). To assess this further, channels on the probe were grouped based on the broad cortical areas above which they were located (Fig. 1C). A three-way ANOVA revealed that average power spectra showed no significant interaction between frequency, novelty (the initial and final timepoints) and frontal (Fig. 2D), somatosensory (Fig. 2E), parietal (Fig. 2F) or retrosplenial (Fig. 2G) areas (Three-way interaction F(6,23) = 0.49, p = 0.81).

We next aimed to quantify the prevalence of bursts of beta frequency oscillations across the cortical surface of the mouse brain, which have previously been shown to dramatically increase in prevalence during exploration of novel environments^6^. Discrete bursts of beta oscillations were observed across all recording channels (Fig. 3A), and the initial rate of beta bursting was substantially higher during the initial minute of recording, compared to the final beta burst rate. While this appeared to be consistent across all recording sites (Fig. 3B), scalp maps reveal that this effect was most notable in channels above the retrosplenial cortex. Pooling the data into distinct recording areas showed a main effect of both region (F(3, 12) = 4.4, P=0.026) and novelty (F(1, 4) = 12, P=0.027), and an interaction between both factors (F(3, 12) = 7.3, P=0.0049, 2-way repeated measures ANOVA). Furthermore, post-hoc Tukey’s multiple comparisons testing revealed a significant difference between initial and final burst rates in the retrosplenial (Mean ± SEM bursts per minute; Initial: 4.02 ± 0.45, Final: 1.62 ± 0.21, P = 0.00011), and somatosensory areas (Initial: 2.92 ± 0.19, Final: 1.61± 0.16, P = 0.0095), but not in the Frontal (Initial: 2.02 ± 0.22, Final: 1.62 ± 0.21, P = 0.36) or parietal areas (Initial: 2.60 ± 0.27, Final: 1.97 ± 0.45, P = 0.17, Fig. 3C-F). To further highlight the topographic variation between regions, we calculated the ‘novelty index’ for each area (initial beta burst rate divided by the final beta burst rate) and found a significant variation in novelty-associated beta bursting across the cortex (F(3) = 6.04, p = 0.01, One-way repeated measures ANOVA, Fig. 3G), with significant differences between retrosplenial (Novelty index: 2.60 ± 0.4) and frontal (1.30 ± 0.24, P = 0.007, Tukey post-hoc test), and parietal areas(1.67 ± 0.45, P = 0.049, Tukey post-hoc test, Fig. 3G).

**Figure 2:**
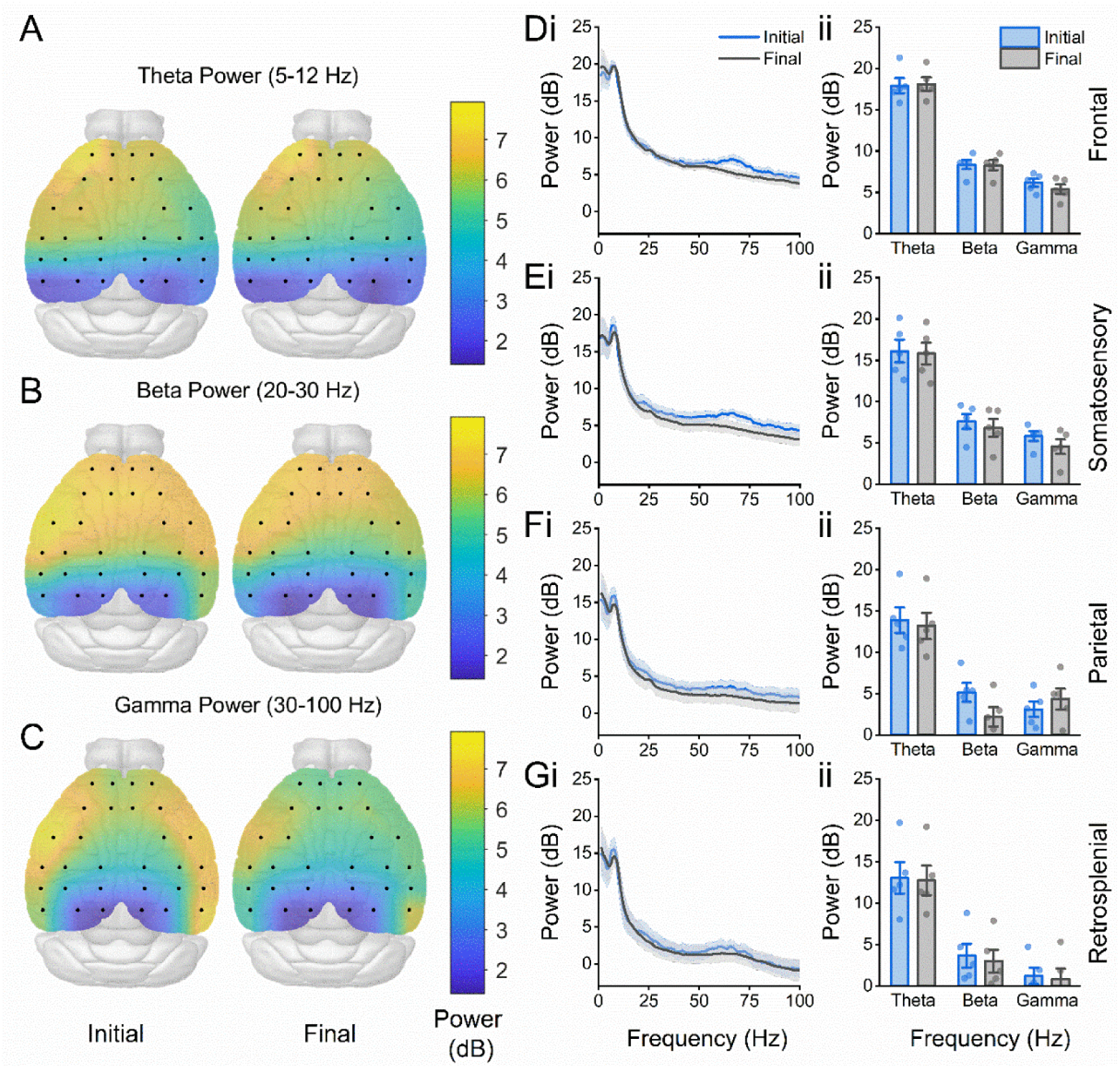
No difference in EEG power spectrum during environmental novelty. **A)** Scalp-map showing total spectral power in the Theta (5-12 Hz) frequency range in the first (left) and final (right) timepoints of the recording session in a novel environment. Equivalent Maps are shown below for total power in the **B)** Beta (20-30 Hz) frequency range and **C)** Gamma (30-100 Hz) frequency range. **D)** Average power spectra (i) and bar charts showing the average power (ii) in frontal **(D)**, somatosensory **(E)**, parietal **(F)** and retrosplenial **(R)** areas during the initial and final stages of the session.

**Figure 3:**
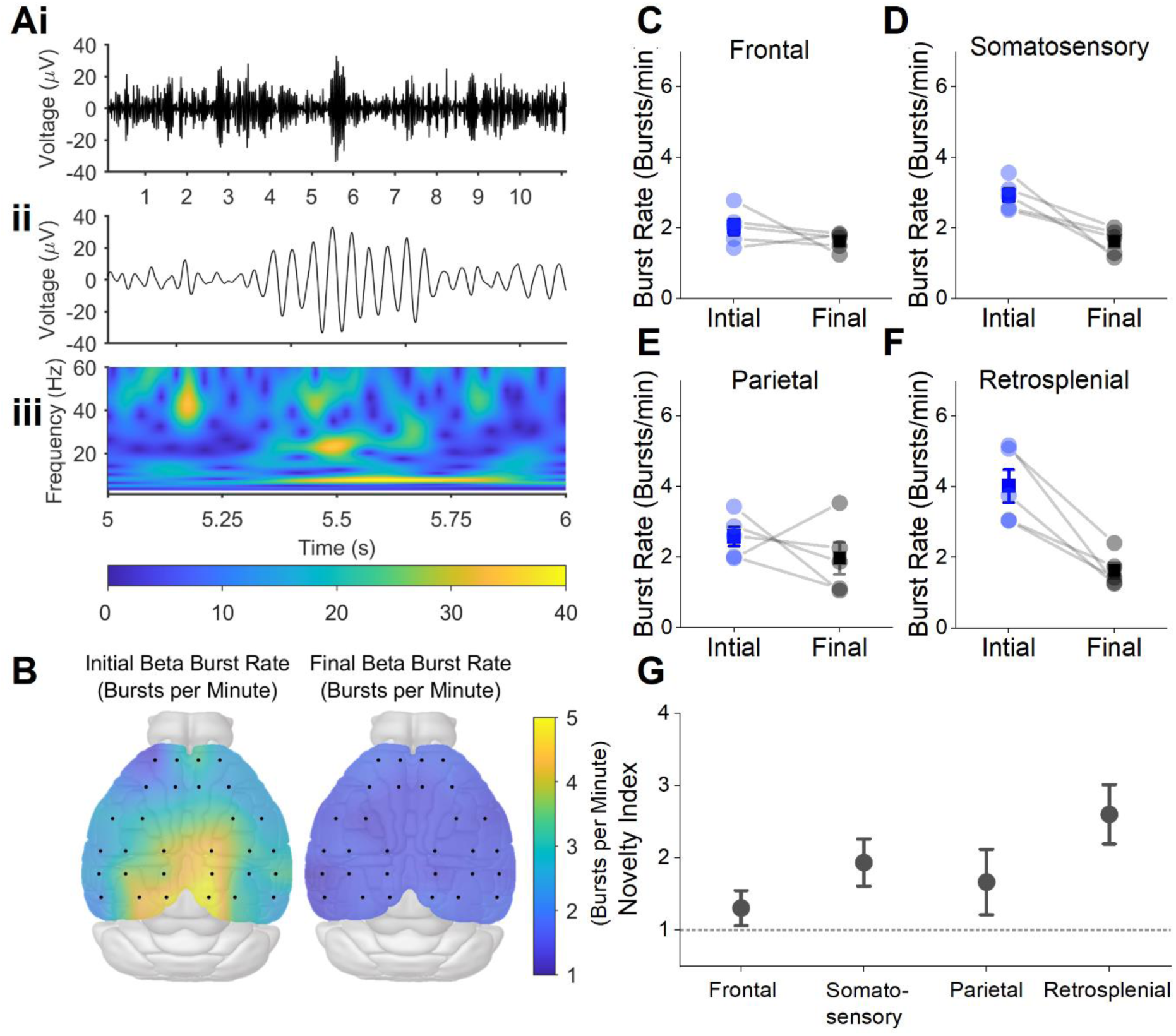
Novelty-associated beta bursting across the mouse EEG. **A)** Example signal from a retrosplenial area recording channel, filtered in the beta frequency range (20-30 Hz) that shows discrete bursts of beta oscillations occurring over 10 seconds **(i),** with an expanded trace showing a single burst **(ii)** and the wavelet spectrogram of this data demonstrating that this event is centred in the beta frequency band **(iii). B)** Scalp maps showing the average beta bursting rate for each electrode in initial (left) and final (right) recording periods, with highest rates of beta bursting occurring in channels above the retrosplenial cortex. Average beta burst rates (mean ± SEM) during the initial and final periods are shown on the right for frontal **(C)**, somatosensory **(D)**, parietal **(E)** and retrosplenial **(F)** channels. **G)** Novelty index (initial burst rate divided by final) for each of the cortical areas with dotted line showing novelty index = 1, which would indicate no effect of novelty on burst rate.

Since spectral analysis revealed some degree of variation in oscillatory power between cortical regions, we next investigated whether this may, at least in part, account for topographical variations in beta bursting. The magnitude of beta bursts followed the overall magnitude of oscillatory activity and varied significantly between cortical areas (F(3) = 18.82, P = 7.8E-5, one-way repeated measures ANOVA, Fig. 4B). However, this was not the case for other burst properties: Total number of beta bursts detected during the recording session was consistent between all cortical areas (F(3) =0.57, P = 0.64, one-way repeated measured ANOVA, Fig. 4A), as was the average duration of beta bursts (F(3) =0.60, P = 0.63, one-way repeated measured ANOVA, Fig. 4C). This suggests firstly, that while beta bursts can be recorded at similar levels across the dorsal cortical surface, novelty-associated changes in beta bursting are specific to certain regions, and secondly, that topographical variation in beta bursting activity across the cortical surface is not merely due to topographical variation in beta burst magnitude.

**Figure 4:**
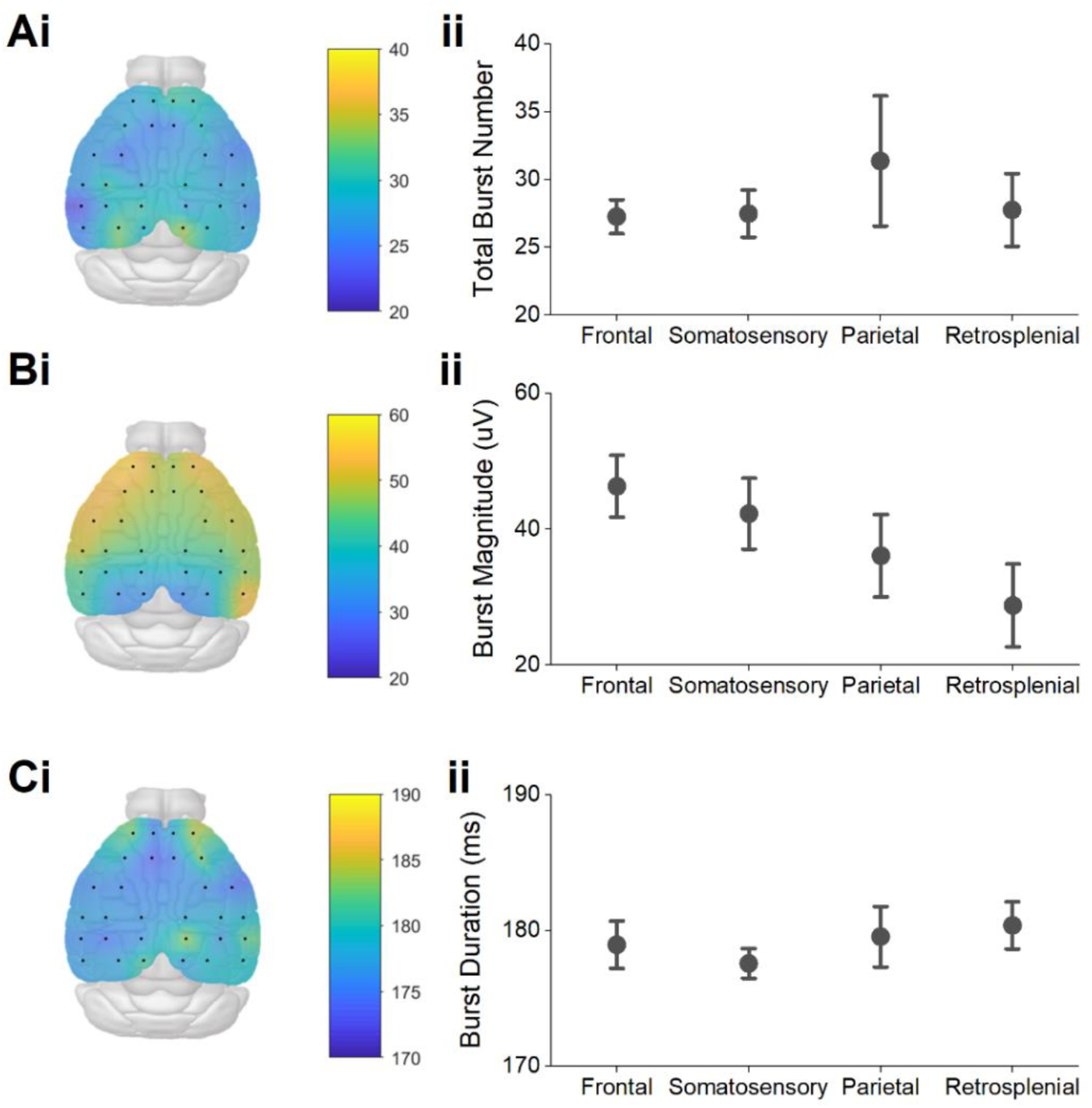
Beta burst properties across the cortical surface. **A)** Scalp map showing the total number of beta bursts detected throughout the whole session at each channel (**i)** and averaged across each cortical area**(ii). B)** Scalp map showing the average beta burst magnitude across the whole session at each channel **(i)** and averaged across each cortical area**(ii). C)** Scalp map showing the average beta burst duration across the whole session at each channel **(i)** and averaged across each cortical area**(ii)** (mean ± SEM).

Since these data suggests that the retrosplenial cortex is highly active during exploration of novel environments, we next sought to quantify the connectivity between this region and the rest of the cortex during the same behaviourally relevant stages of the session. We calculated the amplitude envelope correlation (AEC) between the retrosplenial channels and each of the other channels for data filtered in the theta, beta and gamma frequency ranges (Fig. 5.Ai-iii). There were no significant differences in AEC between the initial and final stages of the session at the single-channel level in the theta frequency range (Fig. 5Ai; within-cluster *U_z_*=8.63, p=0.0938) or in the gamma frequency range (Fig. 5Aiii; within- cluster *U_z_* =30.47, p=0.0625). However, almost all channels showed higher connectivity in the beta frequency range during the initial stage of the session (Fig. 5Aii; within-cluster *U_z_*=26.70, p=0.0313). Moreover, all frequency ranges showed a trend towards higher average RSA-connectivity during the initial stage of the session, but was only significant for theta (Fig. 5Ci; *U_z_*=1.89, p=0.0295) and beta (Fig. 5Cii; *U_z_*=1.89, p=0.0295) frequency ranges, and not for the gamma range (Fig. 5Ciii; *U_z_*=1.62, p=0.0528).

**Figure 5:**
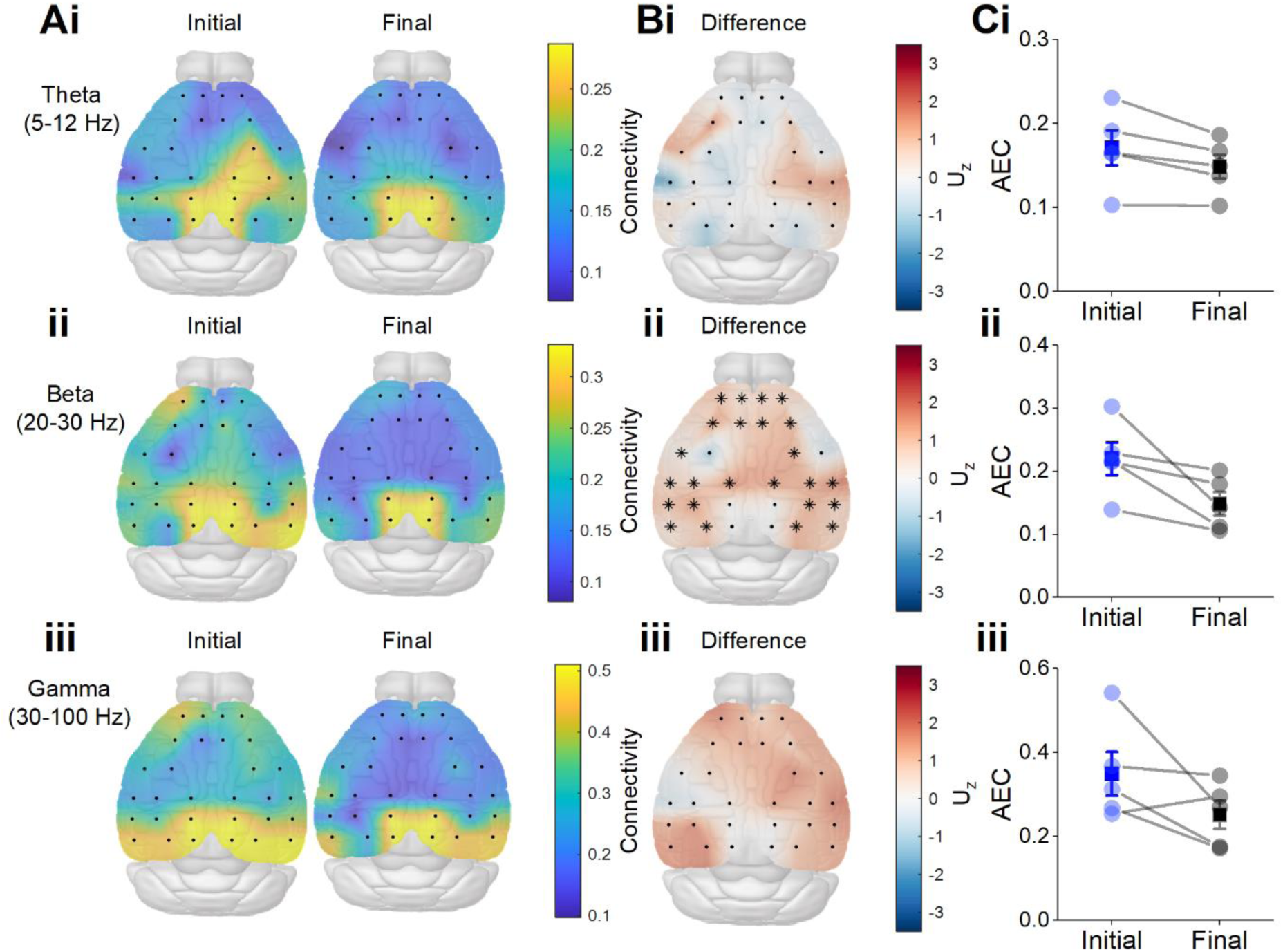
Retrosplenial connectivity during novel recording session. **A)** Scalp maps showing amplitude envelope correlation (AEC) between retrosplenial channels and each of the other channels for data filtered in the theta (**i**), beta (**ii**) and gamma (**iii**) frequency ranges. **B)** Scalp map showing the effect size at each electrode. Asterisks mark significant electrodes using a cluster permutation test for theta (**i**), beta (**ii**) and gamma (**iii**) frequency ranges. **C)** Average retrosplenial-AEC across all channels for each mouse during the initial and final stages of the session for theta (**i**), beta (**ii**) and gamma (**iii**) frequency ranges.

Finally, we asked if changes to connectivity during the initial exploration of a novel environment were specific to the retrosplenial area. Interestingly, when comparing the connectivity differences between each pair of channels, we found that a large proportion of comparisons revealed significant interactions between the initial and final parts of the recording. This was true in both the theta and beta frequency ranges, for which there was a significantly higher AEC across a large portion of edges connecting all channels during the initial stage of the session, suggesting a global increase in AEC (Fig. 6A-B; theta band within-cluster *U_z_* =492.97, p=0.0313; beta band within-cluster *U_z_* =622.96, p=0.0313). No significant differences were found on an edge-wise basis in the gamma range (Fig. 6C; within-cluster *U_z_*=875.38, p=0.0625). As with RSA-seed-based connectivity, in the channel-to-channel connectome the average connection strength (ACS; i.e. the average value of AEC across the whole network) showed trends towards higher global connectivity in all bands, which was significant in the theta (Fig. 6D; *U_z_*=1.89, p=0.0295) and beta (Fig. 6E; *U_z_*=1.89, p=0.0295) bands, but did not reach the threshold for significance in the gamma range (Fig. 6F; *U_z_*=1.62, p=0.0528).

**Figure 6:**
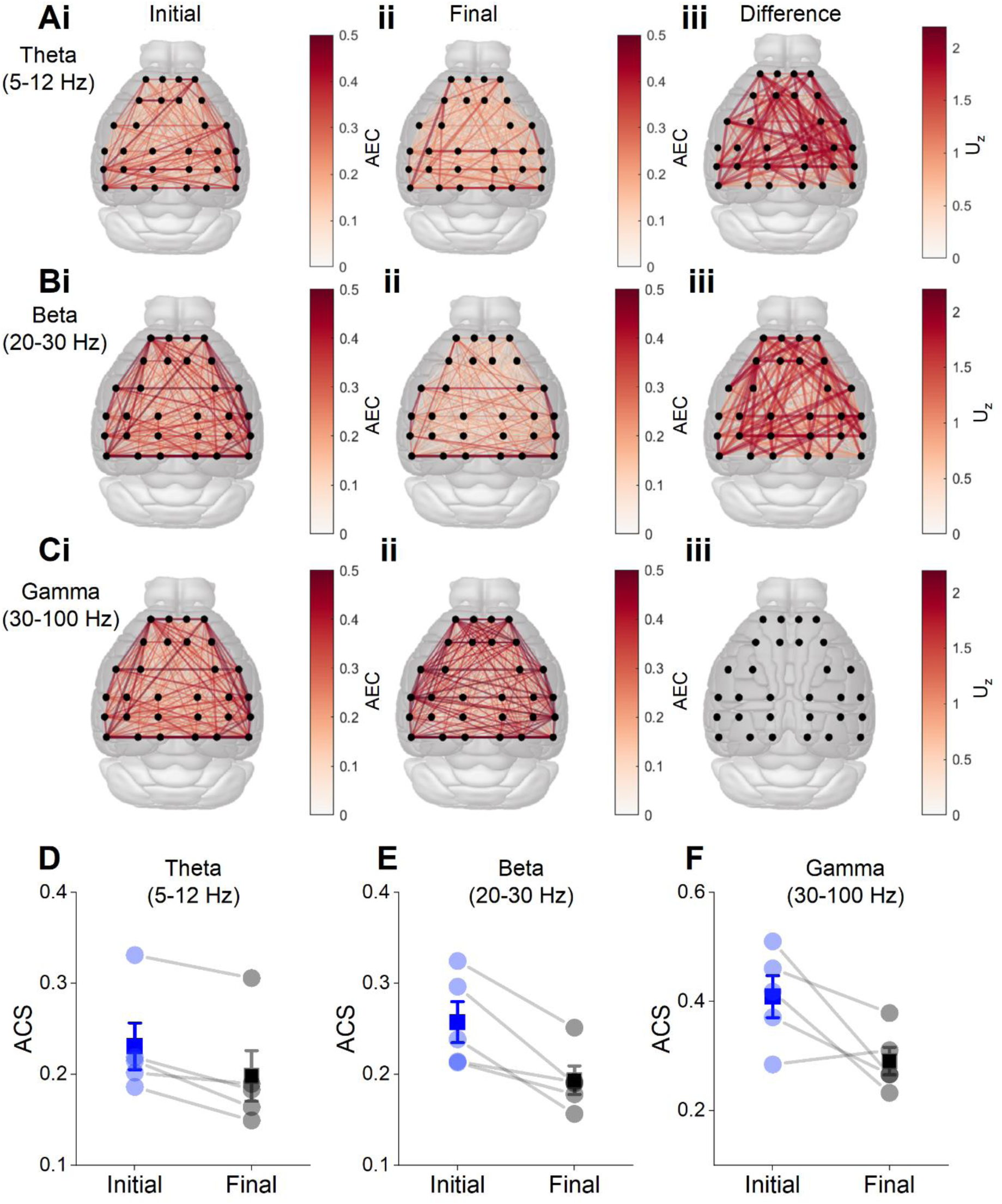
Connectivity across all EEG channels during exploration of novel environment. **Ai-Ci)** Average edge-wise amplitude envelope correlations in the initial minute of exploration in different frequency bands. **Aii-Cii)** Average edge-wise amplitude envelope correlations in the final minute. **Aiii-Ciii)** Difference in connectivity, quantified by the z-approximation of the Mann-Whitney U-statistic. Only edges belonging to significant clusters in the cluster-permutation test are shown. Below: average connection strength (ACS), the average value of AEC across the whole network, showing overall connectivity in the theta (**D**), beta (**E**) and gamma (**F**) frequency ranges.

## Discussion

The aim of this study was to investigate how oscillatory activity changes in response to contextual novelty across the dorsal mouse neocortex, and to determine whether beta bursting activity is a consistent feature of oscillatory activity across the cortex. Beta bursting has been described in numerous cortical regions including the retrosplenial cortex^6^, motor cortex^11^ and somatosensory cortex^17^. Power spectral analysis was performed to investigate oscillatory activity across a range of frequency bands. To investigate the effect of novelty, spectral power was compared between the first minute and final 10 minutes of the session, as previous work has shown that oscillatory responses to contextual novelty peak within the first minute after exposure to the environment and rapidly diminish^1,6,18^. Power spectra from these EEG recordings reveal a broadband attenuation of spectral power compared to power spectra from LFP recordings. Furthermore, heat maps of power in each frequency band reveal a gradient in power across the cortex, with higher power at the rostral end of the cortex, and lower power at the caudal end. The cause of this gradient is unclear, and while it is possible that this is due to the design of these probes, and suggestive of weaker adhesion of channels close to the connector, a similar gradient has been shown previously despite using different probes with the connector at the opposite end^19,20^.

We were able to detect beta bursts across the whole cortex, and the total number of beta bursts detected was generally consistent across all cortical areas. This result supports previous studies demonstrating transient beta oscillations in a range of cortical areas and supports the idea that beta oscillations are a broad cortical phenomenon which underlie a variety of functions depending on their location^11,14,17^.

To investigate the relationship between contextual novelty and beta bursting across the cortex, we compared the rate of beta bursting during the first minute and last 10 minutes of the session. The rate of beta bursting appeared to be higher overall during the novel part of the session compared to the familiar part of the session, especially in the retrosplenial cortex, with an approximate 3-fold increase in the rate of beta bursting in the retrosplenial cortex during novelty. It is important to note that while similar numbers of beta bursts were detected overall in the retrosplenial cortex, the effect of novelty of beta bursting was far more pronounced in the previous LFP studies^6^. These results support the assertation that cortical beta oscillations appear as transient bursts, which may not be apparent in averaged data^6,17^. These data indicate that beta bursting is ubiquitous across the surface of the cortex, but that contextual novelty-associated beta bursting is relatively specific to the retrosplenial cortex and adjacent areas, suggesting that the role of beta bursts in this brain region is related to the processing of contextual information.

As we have shown, beta bursts can be detected across the cortex, it was therefore of interest to investigate the characteristics of these beta bursts, to determine the degree of similarity between beta bursts in the retrosplenial cortex, and those in other cortical regions. There was a significant effect of cortical area on beta burst magnitude, with beta bursts in the retrosplenial cortex being significantly smaller in magnitude than their frontal counterparts. This decrease in beta burst magnitude from rostral to caudal channels mirrored the gradient in spectral power seen earlier, and as before it is unclear whether the cause of this is a true biological gradient, or a technical consideration. The striking consistency of beta burst characteristics across the cortex suggests that the mechanisms underlying the generation of beta bursts are generally similar between different cortical areas, despite vast differences in cytoarchitecture, anatomical connectivity and function between these regions. This supports the idea that beta bursts are generated locally within the cortex and suggests that it is changes in the rate of beta bursting that supports their varying functions within different cortical regions^17^.

We hypothesised that novelty–associated beta bursting would be accompanied by novelty-induced hyperconnectivity. Our results showed a trend towards higher connectivity during contextual novelty across all frequencies, which was significant in the theta and beta bands. Novelty-associated beta bursting was predominantly associated with the retrosplenial cortex, so we started with a retrosplenial seed-based connectivity approach to examine this association. We showed that retrosplenial connectivity was higher during contextual novelty across a large area of the cortex, including frontal, parietal, and somatosensory channels, potentially supporting the hypothesis that beta bursting during novelty provides transient epochs of effective communication across the cortex. However, we found that these changes in connectivity were not specific to retrosplenial connectivity but were identified across the whole network. While novelty associated beta bursting was also demonstrated across the whole cortex, the novelty index was far higher for retrosplenial channels than other channels, an effect which was not reproduced in the connectivity analysis. Hence, it is presently unclear the degree to which the localization of novelty-associated connectivity may be associated with localization of beta bursting and should be the target of future work. The exploration of a novel environment is likely to elicit the activity of numerous, multidimensional networks across a range of oscillatory frequencies^5^ and the complexity of such a system is difficult to quantify with individual metrics.

While we have worked under the hypothesis that connectivity is associated with beta bursting, our methods establish no causality, and the opposite may be true. Additionally, spurious connectivity may be induced via bursting via volume conduction. Since beta bursts are typically high amplitude events, as their field spreads to all electrodes this could potentially induce increased estimates of sensor-level connectivity. However, the use of orthogonalization prior to amplitude envelope computation should in theory eliminate this effect^21^, supporting a physiological association between novelty-associated connectivity and novelty-induced beta bursting.

One limitation to these data is the low spatial resolution of EEG recordings. Due to the spatial filtering properties of the skull, these recordings have far less spatial specificity than depth recordings^22^, therefore in order to account for this, we have referred to these broad groups of channels as “cortical areas” rather than regions. Without source localisation techniques such as those commonly used in human EEG or magnetoencephalography (MEG) studies, it is not possible to determine the specific sources of these oscillations. Additionally, larger cortical areas such as the retrosplenial cortex may therefore have an outsized effect on EEG local field potentials. Additionally, this study focused only on a single novel recording session, and further changes may occur as the animal is repeatedly exposed to the same environment, although previous work by us into novelty-associated beta bursting in the retrosplenial cortex suggests that oscillatory activity during the first minute of exposure to a novel environment is most prominent. Finally, due to the relatively small sample size of this study, while large magnitude changes could be observed, smaller effects may have been missed.

In conclusion, we have demonstrated that beta bursts can be detected across the cortex, that beta burst characteristics are highly consistent between cortical areas, but that novelty–associated changes to beta oscillations are more prominent in retrosplenial adjacent channels. This is accompanied by greater connectivity in the beta frequency range between the retrosplenial and the rest of the cortex. Overall, the cortex is highly modulated by contextual novelty, which may be further investigated in future experiments. These results provide valuable insights into the nature and spatial distribution of beta bursts across the cortex and support our hypothesised role of beta bursting as a means to form cortical representations of context.

## Methods

All procedures were carried out in accordance with the UK Animal (Scientific Procedures) Act 1986 and were approved by the University of Exeter Animal Welfare and Ethical Review Body.

### Animals

Five C57/BL6 mice were bred at the University of Exeter and housed on a 12-hour light/dark cycle. Access to food and water was provided ad libitum. All animals were kept on a 12-hour light/dark cycle, with the light/dark cycle matching the normal daylight/night-time cycle. Mice were group housed prior to surgery, and single housed post-surgery, to prevent damage to the surgical implants.

### Surgery

Mice were fitted with 30-channel EEG probes (NeuroNexus Technologies, MouseEEG), which were fixed to the surface of the skull. Mice were anaesthetised using isoflurane and fixed into a stereotaxic frame. An incision was made along the midline of the scalp, and the skull was revealed. A small droplet of saline was placed around the centre of the skull, upon which the probe was placed and aligned with relation to bregma. Once in place, dental cement (RelyX Unicem, 3M) was applied to the gaps between the rows of electrodes, and around the outside of the probe to hold it in place. Holes were drilled in the frontal bones, and occipital bones, and support screws (Antrin Miniature Specialties) were fitted. The ground wire from the probe was attached to a screw overlying the cerebellum using silver wire. The connector for the probe was manipulated and fixed in position above the probe, and the entire implant was covered with dental cement for support. Throughout surgery, body temperature was monitored with a rectal probe and regulated by a feedback-controlled heat mat.

### Data acquisition

Animals were given at least 1 week of post-operative recovery before the recording session. EEG signals were recorded at 30 kHz using a OpenEphys (open-ephys.org) acquisition board connected to a RHD 32-channel headstage (Intan Technologies) and referenced to a dedicated reference electrode situated above the cerebellum (Fig. 1B). The headstage and tether were counterbalanced using a moveable, weighted arm to allow for the maximum freedom of movement. Two light-emitting diodes (LEDs) on the headstage and an overhead video camera (Logitech HD Pro Webcam C920, Logitech) were used to continuously track the animal’s location using Bonsai (bonsai-rx.org).

Each animal underwent a single 15-minute-long recording session in a novel arena which they were allowed to freely explore. This novel environment was a high sided square arena with black and white stripes. As shown previously, the most notable neurophysiological responses to novelty occur in the initial minutes after exposure to a novel environment, therefore we theorised that a robust neurophysiological response to novelty could be elicited by a single, brief session. After this 15-minute session, each animal was returned to their home cage. To reduce the stress, animals were acclimatised 3 days prior to the start of the experiment, in which they were tethered and recorded from while in their home cage, however none of the animals were placed in the arena prior to their designated recording session.

### Data Analysis

EEG-style surface probes allow sampling of electrophysiological data from across the entire surface of the cortex, however, these recordings have far less spatial specificity than depth recordings^23^. To stratify analysis in this study, channels on the probe were grouped based on the broad cortical areas above which they were located. These were Frontal (F), Somatosensory (S), Parietal (P), and Retrosplenial (R). Of the 30 channels on the probe, 4 electrodes were excluded from this analysis, as they were found at the borders between multiple areas (Fig. 1C).

Analysis was performed on each channel individually and averaged across channels within the same brain region for much of the data. For the construction of heat maps, the built-in MATLAB function scatteredInterpolant was used to assign each value to the coordinate of its channel with relation to bregma and interpolate between these scattered datapoints. Natural neighbour interpolation was used to interpolate between datapoints within the convex hull, but no extrapolation was performed outside the convex hull using scatteredInterpolant. Nearest neighbour extrapolation using scatteredInterpolant treats the nearest neighbour as the nearest true datapoint, rather than the outside edge of the convex hull, resulting in edge effects. Instead, we performed a “skirt” extrapolation, by which nearest neighbour extrapolation was instead performed on the outside edge of the convex hull, resulting in far superior heat maps. Heat maps were overlaid on images taken from Allen’s Brain Explorer^24^.

### Power Spectral Analysis

LFPs were downsampled to 1 kHz for spectral analysis and de-trended, to remove any slow linear drift of the baseline that may occur across the session. Multi-taper spectral analysis was performed using the mtspecgramc function from the Chronux toolbox^25^ (http://chronux.org/), with a time-bandwidth product of 2 (1 second x 2 Hz) and 3 tapers, resulting in some smoothing of resulting spectra. The mtspecgramc function generates a power spectrogram by generating multiple power spectra for short segments of time series data, using a moving window; in our case with the window size of 1 s with no overlap. These spectrograms were then logged to the base 10, and multiplied by 10, to correct for the tendency of spectral power to decrease with a 1/f distribution. These individual spectra were averaged across the first minute (initial) and the last 10 minutes (final) 10 minutes of the recording session. Spectral data from 47 to 53 Hz and from 97 to 103 Hz, which incorporates line frequency noise (50 Hz), and the 100 Hz harmonic were removed and linearly interpolated. The power of each frequency band was calculated as the mean power in each of the following frequency ranges: theta (5-12 Hz), beta (20- 30 Hz) and gamma (30-100 Hz). Wavelet analysis was performed using the cwt function in MATLAB, with the Morlet wavelet with equal variance and time and frequency. The scale to frequency conversions was set by the sampling rate of 1 kHz.

### Beta Burst Detection

Signals were downsampled to 1 kHz for beta burst detection. as in ^6^ and bandpass filtered between 20- 30 Hz to isolate the beta frequency band, using a Butterworth IIR filter with an order of 2. The amplitude and phase of this beta signal were calculated as the real and imaginary components of the Hilbert transform, respectively. The amplitude was z-scored, to give the instantaneous standard deviation of the beta signal amplitude from the mean. Epochs of the signal where this z-score exceeded 2 standard deviations from the mean amplitude were detected, as were the corresponding “edges” of these epochs, where the signal magnitude surpassed 1 standard deviation either side of the 2 standard deviation threshold. This was done to capture the time-course of these high beta amplitude epochs. Events that did not persist longer than a minimum duration of 150 ms (i.e., fewer than 3 oscillation cycles) were discarded. Furthermore, due to the sensitivity of this method to large, amplitude noise artefacts, any event whose peak amplitude exceeded three scaled median absolute deviations from the median of the events detected in that session were discarded. These remaining events were then considered beta-bursts. The duration and peak magnitude of each burst was calculated, as well as the distribution and total number of bursts in the session.

### Connectivity analysis

To test whether novel environments alter the functional connectivity profile, amplitude envelope connectivity (AEC) analysis was performed on EEG data recorded from the initial and final minute of the task following an established pipeline with high test-retest reliability^26^. The data was filtered into a narrow band (theta 5-12 Hz, beta 20-30 Hz, or gamma 30-100 Hz) and orthogonalized using symmetric multivariate orthogonalization to reduce the effects of common sources or spatial leakage due to volume conduction^21^. The amplitude envelope of the orthogonalized data was then calculated via the Hilbert transform, low-pass filtered at 2 Hz and down-sampled to 1 Hz. Subsequently, for each 1-second epoch (i.e., each sample in the down-sampled amplitude envelope), the average movement speed was calculated. Epochs in which the mouse was not moving (movement speed < 2cm/s) were rejected. Since there may be a different number of moving epochs in the initial and final minute, we rejected a subset of epochs in the minute with the largest number of epochs such that (A) both minutes had the same number of epochs and (B) the first and last minute were optimally speed matched. Finally, the samples in the amplitude envelope corresponding to the selected epochs were taken forward for between-channel correlation analysis, forming a 30-by-30 matrix of amplitude envelope correlation coefficients.

In addition to channel-to-channel connectivity, we performed seed-based connectivity analysis using the RSA channels as seed. To calculate RSA-connectivity patterns, for each channel the correlation coefficients with the four RSA channels were averaged, resulting in each non-RSA channel having a single value of RSA connectivity.

### Statistics

For the connectivity analysis, since AEC values are bounded between zero and one and hence not typically normally distributed, we used the Mann-Whitney U-test and its z-statistic ( *U_z_* ) output by Matlab’s “signrank function” (Matlab version R2021a) as a measure of effect size. For the seed-based connectivity analysis, cluster-based permutation testing^27^ was used to identify clusters of electrodes which had significantly different RSA-connectivity in the first and last minute. Cluster-based permutation testing has the advantage that it minimises the problem of multiple hypothesis testing across electrodes. In short, we thresholded *U_z_* values for RSA-connectivity maps to identify clusters of channels/edges with large effect size. For each cluster, we calculated the total within-cluster *U_z_*. We then permuted the values, swapping initial/final labels, and calculated the maximum within-cluster *U_z_* in the permuted data. For a paired test with a sample size of N=5 mice, there are a total of 31 permutations of the data, and hence this was performed for each permutation (as opposed to bootstrap sampling permutations commonly performed for larger sample sizes). The p-value for a cluster was then computed by comparing the empirical within-cluster *U_z_* to the distribution of permuted values, giving a minimum p-value of p=1/31=0.0323. Hence, significance was only achieved if the empirical within-cluster *U_z_* was greater than all permutations. In the channel-to-channel connectivity analysis, the same cluster-based permutation testing approach was taken to find clusters of significant edges in the network. For comparing the global (average) connectivity strengths between the first and final minutes, the Mann-Whitney U test was used.

## Acknowledgments

CW and TR were funded by an ARUK pump priming award. CW was funded by a University of Exeter and Janssen Pharmaceutica studentship. JB was supported by an ARUK Major Project grant (ARUK- PG2017B-7). This work was generously supported by the Wellcome Trust Institutional Strategic Support Award (WT105618MA). For the purpose of open access, the author has applied a CC BY public copyright licence to any Author Accepted Manuscript version arising from this submission.

## Author contributions

**CW:** Conceptualization, Methodology, Formal analysis, Investigation, Writing - Review & Editing, Visualization. **LT:** Formal analysis, Writing - Review & Editing, Visualization. **MG:** Investigation. **JB**: Conceptualization, Resources, Supervision. **TR:** Conceptualization, Formal analysis, Investigation Writing - Original Draft, Visualization, Supervision, Funding acquisition.

## Data Availability

All data will be made available via The Center for Open Science after publication.

## Conflict of Interest

The authors declare no competing financial interests.

## Notes

### Competing Interest Statement

The authors have declared no competing interest.

